# Chronic central-targeted Interleukin-6 overexpression promotes hippocampal and cortical neuropathology in the Tg2576 mouse model of Alzheimer’s disease

**DOI:** 10.1101/2025.02.06.636831

**Authors:** Carla Canal, Kevin Aguilar, Gemma Comes, Elisenda Sanz, Mercedes Giralt, Paula Sanchis, Juan Hidalgo

## Abstract

Interleukin-6 (IL-6) is a cytokine detected in the brains and peripheral fluids of both Alzheimer’s disease (AD) patients and mouse models, where it colocalizes with amyloid-beta (Aβ) levels and amyloid plaques. Interestingly, IL-6 deficiency ameliorates cognitive deficits and attenuates hippocampal neuroinflammation, whereas astrocyte-targeted IL-6 signaling via its soluble receptor accentuates pathological features in AD mouse models. This finding suggests that central IL-6 overexpression may actively drive disease manifestations. However, because IL-6 also signals through its classical membrane-bound receptor pathway, the overall impact of central IL-6 on Alzheimer’s disease pathophysiology is still not fully elucidated. To explore the contribution of central IL-6 overexpression in modulating AD-related mortality, metabolic, behavioral and neuroinflammatory changes in the hippocampus and cortex, we crossed a transgenic mouse model (Tg2576) of Aβ-driven amyloidosis with mice expressing IL-6 under the Glial Fibrillary Acidic Protein (GFAP) promoter, which predominantly targets astrocytes. Chronic IL-6 overexpression reduced inguinal white adiposity in both males and females and decreased body weight in females. Early behavioral alterations were also observed, along with increased cortical and hippocampal Aβ_42_/Aβ_40_ ratios and gliosis in aged Tg2576 female and male mice. Interestingly, chronic IL-6 overexpression also decreased cortical and hippocampal periplaque astrocytosis and microgliosis, suggesting a heterogeneous response of astrocytes and microglia to IL-6 overexpression within the primary regions affected by this pathology. Finally, cortical transcriptomic profiling in Tg2576 mice revealed widespread changes in immune, synaptic, and stress response pathways in response to chronic IL-6 overexpression, with cortical neuroinflammatory and neurotransmission-associated gene networks showing sex-dependent differences. Our findings emphasize that chronic central-targeted IL-6 overexpression shapes the cortical and hippocampal molecular landscape underlying amyloidosis in both male and female Tg2576 mice. Thereby, we propose IL-6 as a potential target for future AD therapeutic strategies.

## Introduction

Alzheimer’s disease (AD) stands as the primary cause of dementia among the aging population, with no significant breakthroughs in the discovery of effective disease-modifying treatments despite extensive research efforts over recent decades [1]. The formation of extracellular amyloid-β (Aβ) deposits and intracellular neurofibrillary tangles composed of hyperphosphorylated tau protein are the two main histopathological hallmarks of AD, primarily affecting vulnerable brain regions such as the hippocampus and the cortex. Beyond these canonical features, the disease is characterized by widespread neuronal dysfunction, synaptic loss, vascular compromise, and chronic neuroinflammation [2]. Importantly, neuroinflammation, driven predominantly by glial responses involving microglia and astrocytes and their release of inflammatory mediators, has been implicated in both the initiation and propagation of AD pathology. While acute glial reactivity may exert protective effects by restricting Aβ accumulation and modulating the local microenvironment, chronic glial reactivity establishes a deleterious feedback loop that amplifies Amyloid Precursor Protein (APP) production and processing, sustained Aβ generation, and the severity of disease progression [3–8].

Interleukin-6 (IL-6) is a multifunctional cytokine that sits at the crossroads of immune regulation, metabolic control, and central nervous system homeostasis. Its dual roles, being both pro-inflammatory and anti-inflammatory, make IL-6 a key participant in the landscape of brain health and disease. Elevated IL-6 levels have been reported in the serum, cerebrospinal fluid, and neural tissue of AD patients, often in proximity to amyloid plaques [7,9]. Additionally, IL-6 overexpression has been observed in translational AD mouse models, where it co-localizes with β-amyloid plaques [10]. In particular, it has been described that IL-6 not only promotes *APP* expression and enhances the levels and activity of Beta-site APP Cleaving Enzyme 1 (BACE1), thereby facilitating Aβ production and perpetuating the positive feedback loop that accelerates AD pathology [7,11], but also can promote gliosis and enhance Aβ clearance via phagocytosis [5,12–14].

Central to the dual effects of this cytokine are the three distinct signaling pathways it activates: classical signaling, which requires the membrane-bound IL-6 receptor; trans-signaling, which involves a soluble form of the IL-6 receptor; and cluster signaling, which occurs between dendritic cells and T cells [15,16]. Interestingly, blocking IL-6 trans-signaling in mainly astrocytes rescues AD-induced mortality and improves behavioral traits and molecular signatures in AD mouse models, primarily in females [17], hence highlighting the importance of astrocytic-targeted IL-6 in this disease and suggesting a sex-dependent effect of IL-6 in AD pathology. However, since IL-6 also signals through its classical membrane-bound receptor pathway, the overall impact of the overexpression of this cytokine on Alzheimer’s disease pathophysiology and how it modulates neuroinflammation in cortical and hippocampal amyloid plaques and its surrounding remains incompletely understood.

By breeding Tg2576 transgenic mouse line, which overexpresses human APP with the Swedish KM670/671NL mutation and displays progressive cognitive deficits and amyloid deposition [18–21], with mice expressing IL-6 specifically under Glial Fibrillary Acidic Protein (GFAP) promoter, we uncovered that chronic central-targeted IL-6 overexpression modulates the molecular landscape underlying amyloidosis in both sexes. In particular, chronic IL-6 overexpression increases cortical and hippocampal Aβ_42_/Aβ_40_ ratios and gliosis in the vicinity of amyloid plaques in aged Tg2576. Moreover, it affected cortical signature of Tg2576 mice, differentially affecting cortical neuroinflammatory and neurotransmission-associated gene networks in males and females. Thus, we position central IL-6 as key regulator of amyloid pathology and of the cortical neuroimmune microenvironment in this model.

## 1. Materials and methods

### 1.1. Ethical statements

All experimental protocols were approved by the Ethics Committee on Animal and Human Experimentation (CEEAH) of the Autonomous University of Barcelona (3971 and 9861) and were conducted in accordance with the Spanish legislation and the EU directive (2010/63/UE) on ‘Protection of Animals Used for Experimental and Other Scientific Purposes’.

### 1.2. Animals

Tg2576 mice (Taconic Europe) [18] and GFAP-IL6 mice, which mainly targets astrocytic IL-6 [22], were used for this study. Hemizygous Tg2576 male mice were crossed with hemizygous GFAP-IL6 females to generate four experimental genotypes: WT (*hAPP*^+/+^, Gfap-Il6^+/+^), GFAP-IL6 (*hAPP*^+/+^, Gfap-Il6^Tg/+^), Tg2576 (*hAPP*^Tg/+^, Gfap-Il6^+/+^), and Tg2576/GFAP-IL6 (*hAPP*^Tg/+^, Gfap-Il6^Tg/+^). Genotypes were verified by PCR, as previously described [22–24]. Mice were group-housed with littermates under specific pathogen-free conditions in a temperature-controlled environment (22 ± 2 °C) with a 12-hour light/dark cycle and *ad libitum* access to food and water. Experimental procedures were performed during the light phase. Mice were monitored regularly from weaning until euthanasia at 18 months of age. Both sexes were included in this study. All procedures were approved by the Ethics Committee on Animal and Human Experimentation (CEEAH) of the Autonomous University of Barcelona (3971 and 9861) and conducted in compliance with Spanish legislation and the EU directive (2010/63/UE) on ‘Protection of Animals Used for Experimental and Other Scientific Purposes’.

### 1.3. Behavioral tests

Behavioral testing was conducted at 5-6 months (young) and at 16–17 months (old), akin to methodology previously described [17,25]. Mice were handled daily for five days before testing for habituation. Each test session was preceded by a 30-minute acclimation period in the home cage within the testing room. Tests were conducted on alternate days in the following order: open field (OF), hole board (HB), elevated plus maze (EPM), pole test, and Y-maze (YM) for working and spatial memory. Data collection and analyses were performed under blind conditions. All tests were recorded using an overhead video camera, and data were collected through both *in situ* manual measurements and automatic tracking analysis using EthoVision XT software, version 11 (Noldus).

#### 1.3.1. Open Field, Hole Board, and Elevated Plus Maze tests

Locomotor and exploratory activity were assessed in the OF (36,5 x 56 x 31 cm) over 5 minutes. Total horizontal (ambulatory distance) and vertical (rearing count) activities were recorded. Exploratory behavior was assessed in the HB (40 x 40 x 21,5 cm, four equally spaced bottom holes of 3 cm diameter) over 5 minutes. Time and number of head-dipping events were measured. Anxiety-like behavior was assessed in the EPM (cross-shaped maze consisting of two 20 x 5 cm open arms and two 20 x 5 x 15 cm closed arms and elevated 47 cm above the floor) for 5 minutes. Percentage of entries and time spent in the open arms (OA) of the apparatus (excluding the center) were quantified.

#### 1.3.2. Motor coordination

Motor coordination and balance of mice were evaluated using a vertical pole (30 cm height, 1 cm diameter, rough surface) for 120 seconds. After three habituation trials conducted one week beforehand, mice performed three consecutive trials separated by 10 minutes. The time to descend (maximum duration was scored if the mouse fell down the pole) and the number of failed trials were measured. Motor dysfunction and cerebellar ataxia were further assessed using the Semi-Quantitative Cerebellar Ataxia (SQCA) scale (scoring adapted from [26,27]), scoring hindlimb clasping, ledge walking, and kyphosis (each 0-3; total 0-9) from no dysfunction to severe ataxia. Three trials per component were averaged for final scoring.

#### 1.3.3. Y-Maze tests

Working and spatial reference memory were evaluated using a Y-shaped apparatus (33 x 5 x 15 cm) with external visual cues. For working memory, mice were allowed to freely explore the apparatus in a single 8-min trial; spontaneous alternations (three consecutive entries to three different arms) were calculated as (alternations / [total arm entries – 2] x 100). For spatial reference memory, mice were allowed to freely explore the apparatus for 10 min with one arm closed off (training session). After 1h and 2h retention intervals, the blocked arm was reopened for 5 minutes (trial sessions). Entries and time in the novel arm (NA) of the apparatus were recorded.

### 1.4. Neuropathological analysis

#### 1.4.1. Sample collection

At 18 months, mice were weighed and euthanized by decapitation. Fat depots including interscapular brown adipose tissue (BAT) and subcutaneous inguinal white adipose tissue (WAT) were dissected and weighed. Brains were extracted and the left hemisphere was fixed in 4% paraformaldehyde (PFA) for 24 h at 4 °C for further processing, while the right hemisphere was dissected into cortical (anterior and posterior) and hippocampal regions, snap-frozen in liquid nitrogen, and stored at −80 °C for molecular analysis.

#### 1.4.2. Immunofluorescence (IF)

Fixed hemispheres were embedded in OCT, and sagittal sections (20 μm thick) were cut using a cryostat (Leica CM3050 S). All the procedures in this subsection were performed in free-floating sections. Sections were blocked with 1% BSA/PBS + 0.2% Triton X-100. For amyloid detection, sections underwent 70% formic acid pretreatment (5 min). Primary antibodies (mouse anti-β-amyloid_17-24_ 4G8, Biolegend; rabbit anti-GFAP, Dako; and rabbit anti-Iba1, Wako) were incubated 1:500 overnight in blocking solution, followed by secondary antibodies (goat anti-mouse IgG (H+L) Alexa Fluor™ 488 1:600; Invitrogen A11001; and donkey anti-rabbit IgG (H+L) Alexa Fluor™ 594 1:600; Invitrogen A21207) 1:600 in blocking solution. Slides were mounted and covered with DAPI Fluoromount-G^®^ mounting media (Southern Biotech 0100-20). For quantification, three non-consecutive sections per mouse were selected and imaged using an EVOS™ M5000 Imaging System (Invitrogen) microscope, and images were analyzed using ImageJ software [28]. The sum of intensity of the specific signal (integrated density) was determined for cortex and hippocampus and normalized by the area of the region of interest (Area ROI). The integrated density values for GFAP and Iba1 were quantified within a fixed-width peri plaque annular region and normalized by ROI area (plaque-shaped areas).

#### 1.4.3. Western Blot

Tissue samples were homogenized by sonication in a lysis buffer (50 mM Tris-HCl pH 7.6, 150 mM NaCl, 2mM EDTA, 0,1% Igepal CA 630, 3% SDS, and 2% sodium deoxycholate, supplemented with a protease inhibitor cocktail and phenylmethylsulfonyl fluoride). Total protein was quantified via the Pierce^TM^ BCA Protein Assay Kit (ThermoFisher) according to the manufacturer’s protocol.

Lysates were mixed with NuPAGE™ LDS 4X (Invitrogen) and 2-mercaptoethanol (Sigma-Aldrich) and were heat shock for 5 min at 90 °C. Proteins were separated by electrophoresis on a NuPage^TM^ 4–12% Bis-Tris Midi protein Gel (Invitrogen) and transferred to 0,2 μm nitrocellulose membranes. Membranes were blocked with 5% (w/v) skim milk, incubated with overnight primary antibodies (mouse anti-β-amyloid_1-16_ 6E10 1:2000, 803003 Biolegend; rabbit anti-GFAP 1:40000, Z0334 Dako; and rabbit anti-Iba1 1:3000, 019-19741 Wako) at 4 °C, and with secondary antibodies (polyclonal rabbit anti-mouse Ig/HRP 1:5000, P0260 Dako; and goat anti-rabbit IgG (H+L)-HRP conjugate 1:5000 and 1:10000, 1706515 Bio-Rad). Immunoreactive bands were detected using the Clarity Western ECL Substrate (Bio-Rad), imaged in the Chemidoc Imaging System (Bio-Rad), and analyzed using Image Lab^TM^ 6.0 Software (Bio-Rad). Monoclonal anti-β-Actin (1:5000, Sigma-Aldrich) was used as a loading control following membrane stripping with 2% SDS, Tris HCl, and Tris Base pH 6.7.

#### 1.4.4. Enzyme-Linked ImmunoSorbent Assay (ELISA)

Aβ_40_ and Aβ_42_ concentrations were measured with Amyloid beta 40 and 42 Human ELISA Kits (ThermoFisher) in cortical and hippocampal homogenates (as described in 3.3.2). The protocol was performed according to the manufacturer’s recommendations. The protein concentration of the lysate was used to normalize the data.

#### 1.4.5. Luminex multiplex assay

Tissues were homogenized mechanically (MM-400 mixer mill with ceramic beads) and by sonication in a protein extraction buffer (6,25 mM HEPES, 0,02% Igepal CA 630, 5 mM MgCl_2_, 16,9 μM EDTA pH 8.0 and 1 μM EGTA pH 8.0 mM, supplemented with a protease inhibitor cocktail and phenylmethylsulfonyl fluoride). Homogenates were centrifuged (13500 x g, 5 min) and the supernatant was collected. Total tissue protein quantification was performed with the Pierce^TM^ BCA Protein Assay Kit (ThermoFisher) according to the manufacturer’s protocol.

Cytokines (IL-6 and IL-10) were analyzed using the MILLIPLEX MAP Mouse High Sensitivity T Cell Magnetic Bead Panel (Merck Millipore MHSTCMAG-70K) in a Luminex MAGPIX® Instrument System (powered by Luminex xMAP technology) and analyzed using the xPONENT 4.2 software (Luminex Corporation MAGPIXXPON42). The protocol was performed according to the manufacturer’s recommendations. Measurements below the first calibration standard were scored as zero. Data were normalized to total protein concentration.

### 1.5. Bulk RNA-sequencing

Frozen cortical tissue samples were homogenized mechanically in Buffer RLT supplemented with 1% (v/v) β-mercaptoethanol using a Polytron Homogenizer (Kinematica Inc.). Homogenates were centrifuged (2000 x g, 10 min, 4°C), and RNA was extracted from the resulting supernatants using the RNeasy Mini Kit (#74104, Qiagen) according to manufacturer’s recommendations and including the in-column DNase treatment to eliminate genomic DNA contamination. RNA concentration was determined by Nanodrop (Thermo Fisher Scientific). Four biological replicates per group and sex were used.

RNA-Seq and data processing to generate the matrix was performed at Biomarker Technologies (BMK) GmbH (MuDnster, Germany). Sequencing libraries were generated according to the manufacturer’s standard protocol and subjected to paired-end (PE) sequencing (2 × 150 bp) on the Illumina NovaSeq X platform. An estimated Data Output of ∼20 million PE reads per sample was generated. Raw sequencing data were first subjected to quality control filtering using in-house BMK scripts to remove reads containing adapter sequences, reads with poly-N, and low-quality reads. The quality of the clean data was evaluated by calculating Q20, Q30, GC content, and sequence duplication levels. Clean reads from each sample were mapped to the *Mus musculus* reference genome Mus_musculus.GCF_000001635.27.genome.fa. and then aligned to the specified reference genome using HISAT2 [29]. Subsequently, StringTie [30]was used to assemble transcripts based on the alignment results and gene expression levels were quantified using the FPKM method and normalized for gene length and sequencing depth.

Differential expression analysis was conducted using the DESeq2 package [31], with transcripts considered differentially expressed when exhibited at a fold change (FC) greater than 1.5 and an adjusted p-value (padj) below 0.05. For graphical representation, volcano plots were generated using the VolcaNoseR online tool [32], displaying padj values on a -log10 scale and FC on a log2 scale (Log2FC). Gene ontology (GO) analysis was performed using the WebGestalt online tool [33]. Dotplots depicting top GO biological categories [32], Venn diagrams [33] and heatmaps of selected gene expression profiles for each pair-wise comparison were visualized using R package ggplot2 [34] and Venn diagram [35]. The resulting GO biological process enrichments were then visualized using R package Rtsne [36] and ggplot2 [34]. All plots were produced with standardized parameters for clustering and scaling to ensure comparability across datasets.

### 1.6. Statistics

Data are plotted using GraphPad Prism version 10 and presented as mean ± SEM. Statistical analysis was conducted using SPSS version 23. Survival curves were analyzed using Kaplan-Meier and Log-rank survival tests (four-genotype model). Analyses with two independent variables were performed using Generalized Linear Model (GzLM) with post hoc sequential Bonferroni’s correction. Repeated measures were analyzed via Generalized Estimated Equations (GEE), including genotype as a fixed factor and time as a within-subject variable. For comparisons involving only two independent groups, unpaired t-tests were used. Males and females were analyzed separately, with the exception of the cortical transcriptome analysis in which sex differences were taken into account. Statistical significance was set at *p* ≤ 0.05. Sample size is stated in the text or figure as needed. It is important to note that this study is not powered.

## 2. Results

### 2.1. Chronic central-targeted IL-6 overexpression decreased inguinal adipose tissue in Tg2576 males and females, exacerbating body weight loss in Tg2576 females

To examine the effect of central IL-6 overexpression in Tg2576 mice, we first crossed GFAP-IL6 mice, which express *Il6* under the GFAP promoter, with Tg2576 mice and then we monitored the survival and body weight of their offspring: WT, GFAP-IL6, Tg2576, and Tg2576/GFAP-IL6. Whereas GFAP-IL6 females showed no effect on survival and GFAP-IL6 males exhibited a modest survival benefit compared with WT, survival was significantly reduced in both male and female Tg2576 mice. Central IL-6 overexpression did not accentuate mortality in Tg2576 mice (Fig. 1A).

**Figure 1.**
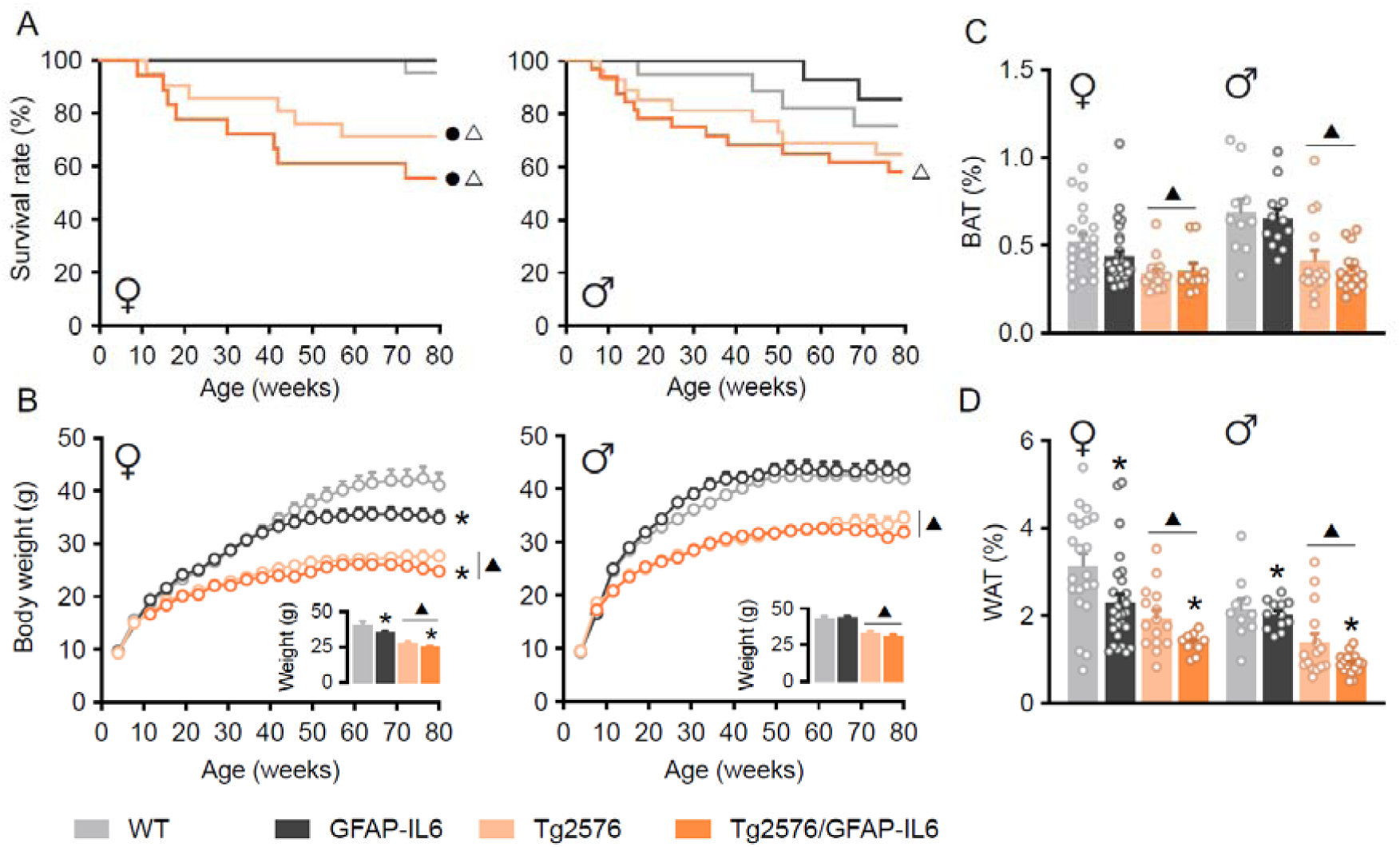
Life span and body composition of offspring from breeding Tg2576 and GFAP-IL6 mice. **A** Kaplan-Meier survival curves for all genotypes, analyzed by log-rank test; ● p ≤ .05 vs WT, △ p ≤ .05 vs GFAP-IL6. **B** Longitudinal body weight and terminal weight at euthanasia *(small)*. **C** Brown adipose tissue relative weight. **D** White adipose tissue relative weight. Data are mean ± SEM; statistics by GEE (B) and GzLM (C-D). ▴ p ≤ .05 Tg2576 effect, **\*** p ≤ .05 IL-6 effect. Female n = 18-30, male n = 20-38 at 3 weeks; female n = 11-30, male n = 14-26 at 80 weeks. BAT, Brown adipose tissue; WAT, White adipose tissue.

Tg2576 animals also consistently maintained lower body weight throughout their lifespan relative to littermate controls. Late-onset weight loss was evident in female mice with central IL-6 overexpression, especially in the GFAP-IL6 genotype, whereas this effect was not observed in males (Fig. 1B). In line with overall body weight trends, both brown (BAT) and white adipose tissue (WAT) weights, normalized to body weight, were significantly reduced in Tg2576 mice. Further, relative WAT fat depots were reduced in IL-6-overexpressing female and male mice compared to littermate controls (Fig. 1C, D). Together, these results reveal that survival and body weight are affected in Tg2576 mice, and that central IL-6 overexpression reduces inguinal adiposity in both Tg2576 female and male mice, with body weight affected only in females.

### 2.2. Chronic central-targeted IL-6 overexpression modulates AD-like behavior parameters in Tg2576 mice from early stages

Since Aβ plaque deposition in Tg2576 mice occurs from 12 months of age [37], we next evaluated whether central-targeted IL-6 overexpression could impact locomotion, exploration, and anxiety in Tg2576 mice before and after Aβ plaque deposition. To this end, WT, GFAP-IL6, Tg2576, and Tg2576/GFAP-IL6 mice were evaluated in various behavioral paradigms at 5-6 and 16-17 months of age (Fig. 2A).

**Figure 2.**
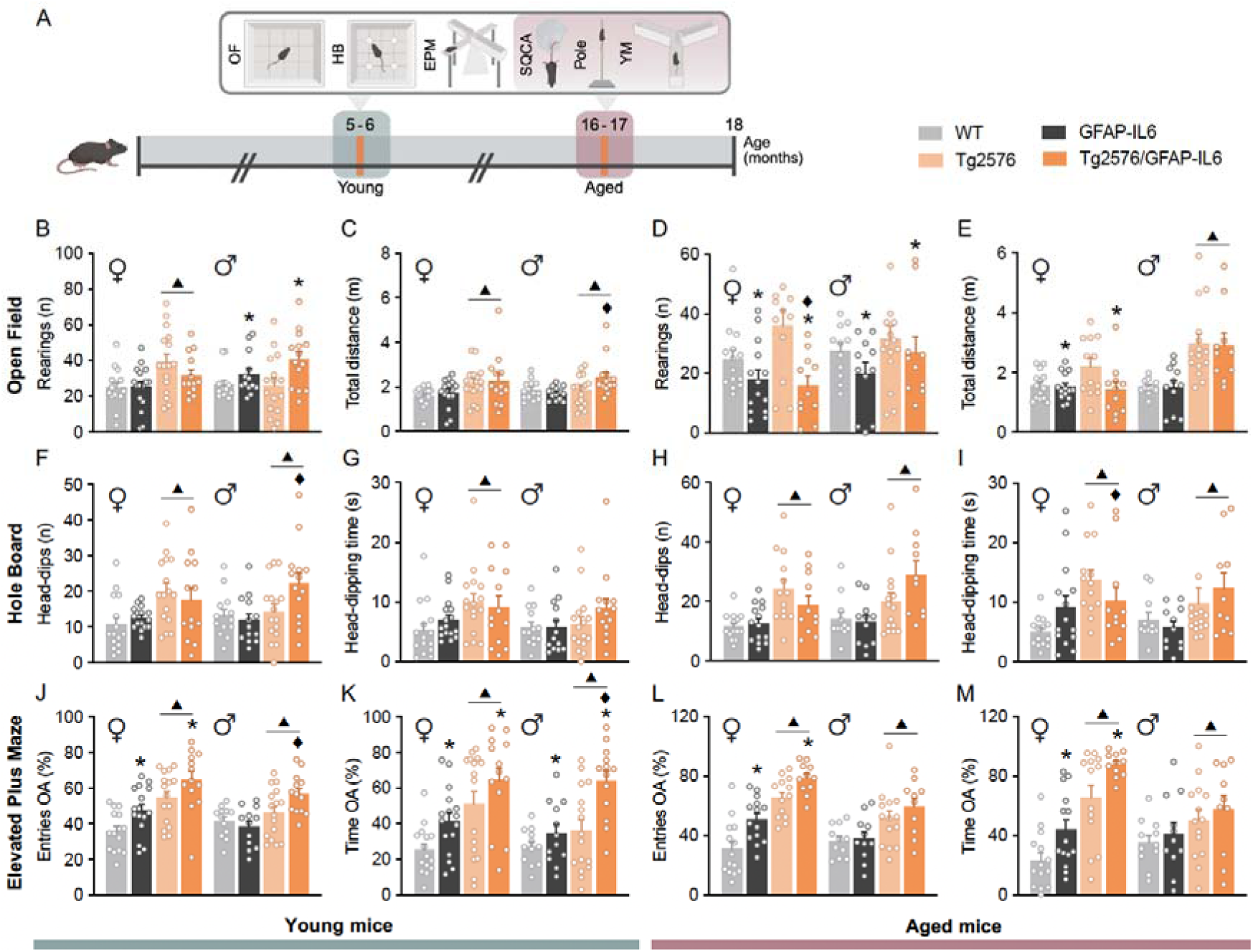
Behavioral assessments in young and aged Tg2576 and GFAP-IL6 mice. **A** Experimental design for behavioral assessments in young (grey) and aged (orange) mice. **B-E** Open field test. Number of rearings (B, D) and total distance travelled (C, E) in young and aged mice, respectively. **F-I** Hole board test. Number (F, H) and duration (G, I) of head-dipping events in young and aged mice, respectively. **J-M** Elevated plus maze test. Percentage of entries (J, L) and percentage of time (K, M) in the open arms in young and aged mice, respectively. Data are mean ± SEM; analyzed by GzLM statistical test. ▴ p ≤ .05 Tg2576 effect, **\*** p ≤ .05 IL-6 effect, ♦L p ≤ .05 interaction between both factors. OA, Open arms.

Young Tg2576 (5-6 months old) mice exhibited hyperactivity in the open field test, marked by increased ambulation (horizontal activity) and rearing behavior (vertical activity) (Fig. 2B, C), enhanced exploratory behavior in the hole board, reflected in more frequent and prolonged head-dipping events (Fig. 2F, G), and decreased anxiety in the elevated plus maze, shown by a higher percentage of entries and time spent in the open arms, with changes being more evident in females (Fig. 2J, K). However, whereas central-targeted IL-6 overexpression attenuated vertical and horizontal activity and did not impact on exploratory behavior in females, it increased vertical and horizontal activity and exploratory behavior in males, particularly Tg2576/GFAP-IL6 male (Fig. 2B, C, F, G). Additionally, both male and female IL-6-overexpressing mice exhibited reduced anxiety-like behavior, especially in Tg2576/GFAP-IL6 mice (Fig. 2J, K).

In aged Tg2576 mice (16-17 months old), these behavioral impairments observed at 5-6 months of age persisted. However, in contrast to the findings in young mice, central-targeted IL-6 overexpression in aged mice reversed hyperactivity in both sexes (Fig. 2D, E) and partially restored exploratory behavior in Tg2576/GFAP-IL6 females while increasing it in males (Fig. 2H, I). Additionally, central-targeted IL-6 overexpression increased both the frequency and duration of entries into the open arms of the elevated plus maze, particularly in Tg2576/GFAP-IL6 female mice, with males showing a similar trend (Fig. 2L, M).

Aged Tg2576 mice experienced significant motor deficits, as evidenced by abnormal SQCA scores, and especially abnormal hindlimb extension and increased kyphosis. Central-targeted IL-6 overexpression worsened hindlimb extension, kyphosis, and motor coordination in the ledge test (Fig. 3A-D), consistent with previous findings [27,38]. Pole test performance also revealed motor impairment in Tg2576 mice, particularly females, as indicated by prolonged descent times, although the number of failed trials remained unchanged. GFAP-IL6 mice also exhibited impaired motor performance, marked by a longer descent time from the pole and a higher number of failed trials. However, central-targeted IL-6 overexpression did not exacerbate the motor impairment in Tg2576 mice (Fig. 3E, F).

**Figure 3.**
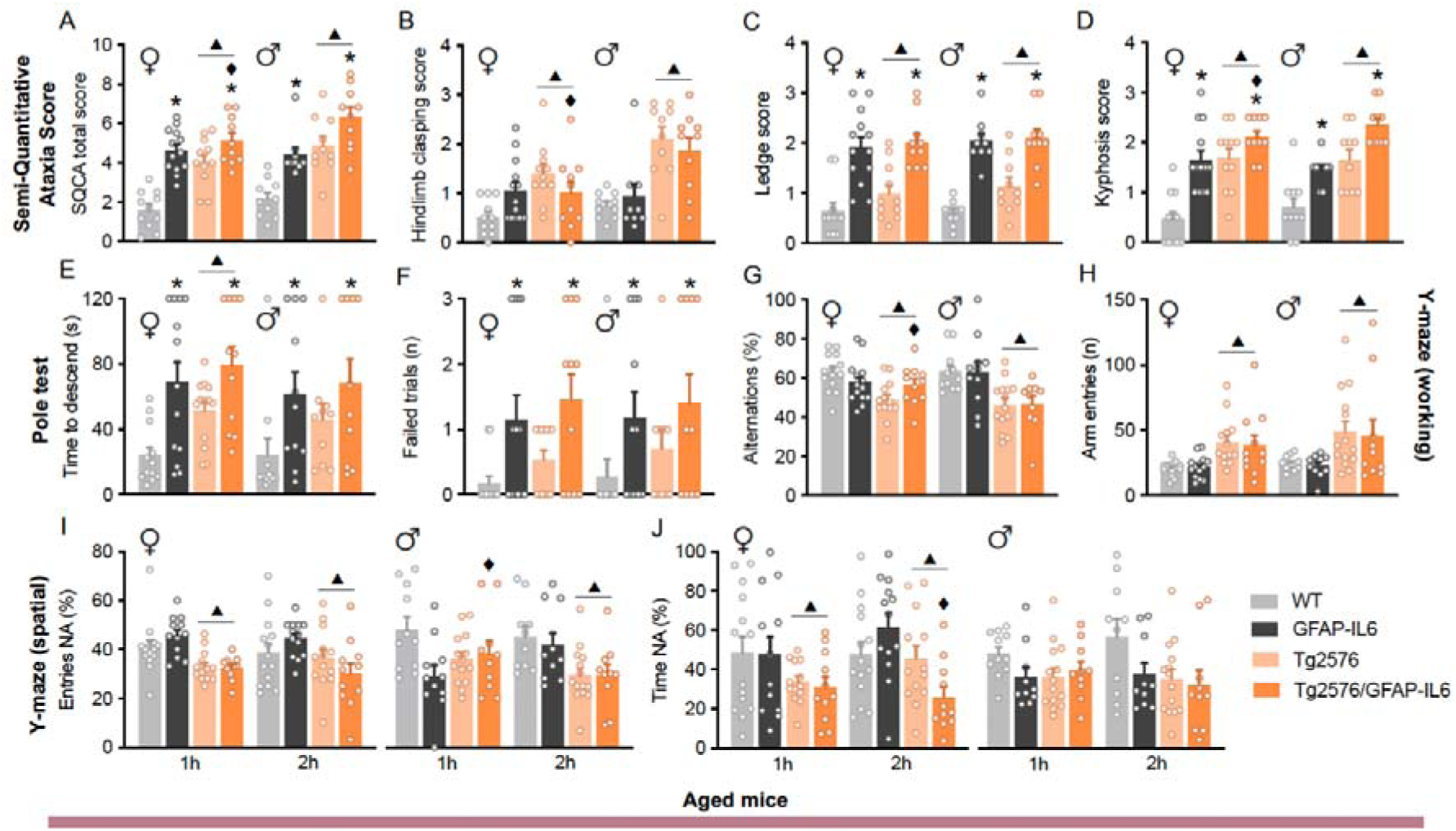
Motor performance and cognition assessment in aged Tg2576 and GFAP-IL6 mice. **A-D** Semi-quantitative cerebellar ataxia scoring. SQCA total (A), hindlimb clasping (B), ledge performance (C), and kyphosis scores (D) in aged mice. **E-F** Pole test. Time to descend (E) and number of failed trials (F) in aged mice. **G-H** Working memory performance in the Y-maze test. Percentage of spontaneous alternations (G) and total arm entries (H) in aged mice. **I-J** Spatial memory performance in the Y-maze test. Number of entries (I) and time spent (J) in the novel arm in aged mice. Data are mean ± SEM; analyzed by GzLM statistical test. ▴ p ≤ .05 Tg2576 effect, **\*** - p ≤ .05 interaction between both factors. SQCA, Semi-quantitative cerebellar ataxia; NA, Novel arm.

Working and spatial memory were impaired in aged Tg2576 mice, with spontaneous alternation and novel arm exploration deficits in the Y-maze (Fig. 3G-J). Whereas central-targeted IL-6 overexpression improved working memory in aged female Tg2576/GFAP-IL6 mice; in male mice, no effects were observed regarding central-targeted IL-6 overexpression. While a higher number of total arm entries was also observed in Tg2576 mice, no effects were related to central-targeted IL-6 overexpression (Fig. 3G, H). Central-targeted IL-6 overexpression had opposite effects in spatial learning and memory in females, as it improved memory in GFAP-IL6 mice while worsening it in Tg2576/GFAP-IL6 mice, particularly during the test performed 2 hours after training. Conversely, GFAP-IL6 male mice displayed worsened spatial memory (Fig. 3I, J).

Collectively, these results suggest that central-targeted IL-6 overexpression partially restores exploratory and working memory deficits while exacerbating motor impairments and spatial memory in aged Tg2576 females. Overall, these findings reveal that central-targeted IL-6 overexpression regulates behavior in Tg2576 mice in an age- and sex-dependent manner.

### 2.3. Chronic central-targeted IL-6 overexpression accentuates cortical and hippocampal Aβ_42_ / Aβ_40_ratios in aged Tg2576 mice

Given that AD-like pathology in Tg2576 mice manifests in the hippocampus and subsequently extends to cortical regions in aging [17], we next evaluated the influence of central-targeted IL-6 overexpression on underlying molecular mechanism of AD-like disease at 18 months of age. As expected, the brains of Tg2576 mice displayed significant Aβ pathology and progressive development of diffuse and senile plaque resulting from hAPP overexpression and processing (Fig. 4A). Quantitative analysis did not reveal differences in amyloid plaque burden in Tg2576/GFAP-IL6 mice compared to Tg2576 littermates (Fig. 4B, C). Both cortical and hippocampal hAPP holoprotein and total Aβ peptide levels were elevated in Tg2576 mice compared to WT controls (Fig. 4D, Supplementary Fig. 1). Although central-targeted IL-6 overexpression did not affect hAPP holoprotein levels, it reduced total Aβ peptide levels in the hippocampus of both male and female mice, and in the cortex of females only (Fig. 4E-H). Further analysis of Aβ species demonstrated that central-targeted IL-6 overexpression selectively decreased Aβ_40_ in the hippocampus of both females and males, and in the cortex of males (Fig. 4I, J), whereas increased Aβ_42_, the more pathogenic variant, in the cortex of both sexes and in the hippocampus of males (Fig. 4K, L). This shift resulted in a dramatically higher Aβ_42_/Aβ_40_ ratio in Tg2576/GFAP-IL6 mice compared to Tg2576 mice both in cortex and hippocampus of males and females (Fig. 4M, N). Collectively, these results suggest that overexpression of central-targeted IL-6 accentuates the AD-like neuropathology.

**Figure 4.**
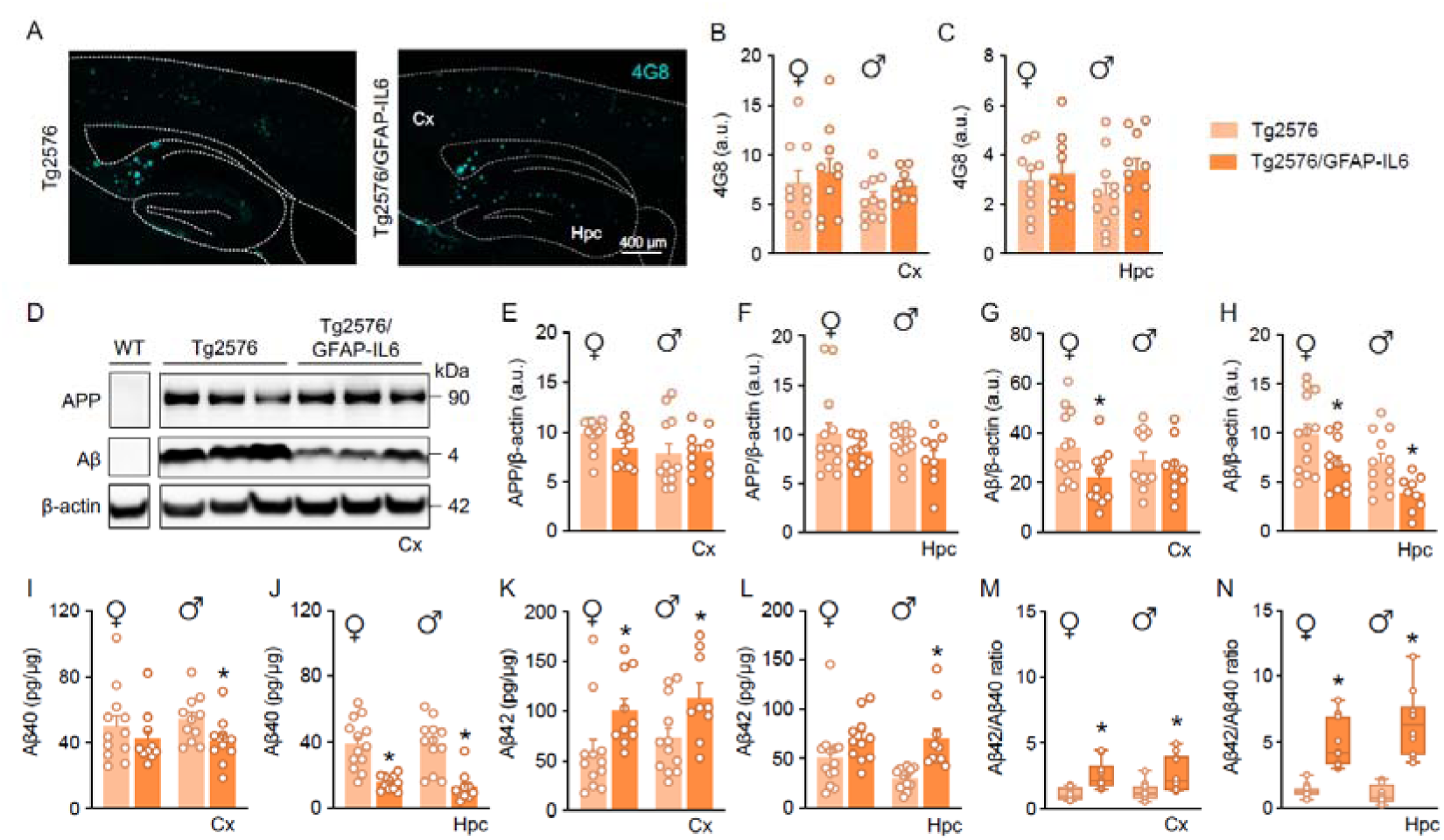
Amyloid pathology in the cortex and hippocampus. **A - C** Representative 4G8 immunofluorescence (A) and cortical and hippocampal quantification of 4G8 levels (B, C). **D - H** Representative cortical 6E10 immunoblot (D) and quantification of APP and Aβ levels relative to β-actin in the cortex (E, G) and the hippocampus (F, H). **I - N** Aβ40 and Aβ42 quantification by ELISA in the cortex (I, K) and hippocampus (J, L), and relative cortical and hippocampal Aβ42/Aβ40 ratio (M, N). Data are mean ± SEM; analyzed by Student’s t-test. **\*** p ≤ .05 IL-6 effect. Cx, cortex; Hpc, hippocampus.

### 2.4. Chronic central-targeted IL-6 overexpression promotes cortical and hippocampal glial reactivity in aged Tg2576 mice

Astrocytes and microglia are major glial cells orchestrating neuroinflammatory responses and play key roles in the pathogenesis of AD. In AD-like neuropathology, reactive glial cells cluster around Aβ deposits and contribute to a complex inflammatory milieu that can influence synaptic function, neuronal survival, and amyloid clearance [3,6,14]. Since both astrocytes and microglia are significant sources and responders to IL-6 [39], we evaluated whether central-targeted IL-6 overexpression affects astrocytic and microglial reactivity after confirming that elevated IL-6 levels were detected in GFAP-IL6 and Tg2576/GFAP-IL6 mice. Notably, IL-6 overexpression was detected both the cortex and, most prominently, in the hippocampus of both male and female without differences in IL-6 levels between wild-type and Tg2576 genotypes in either brain region (Fig. 5A, B).

**Figure 5.**
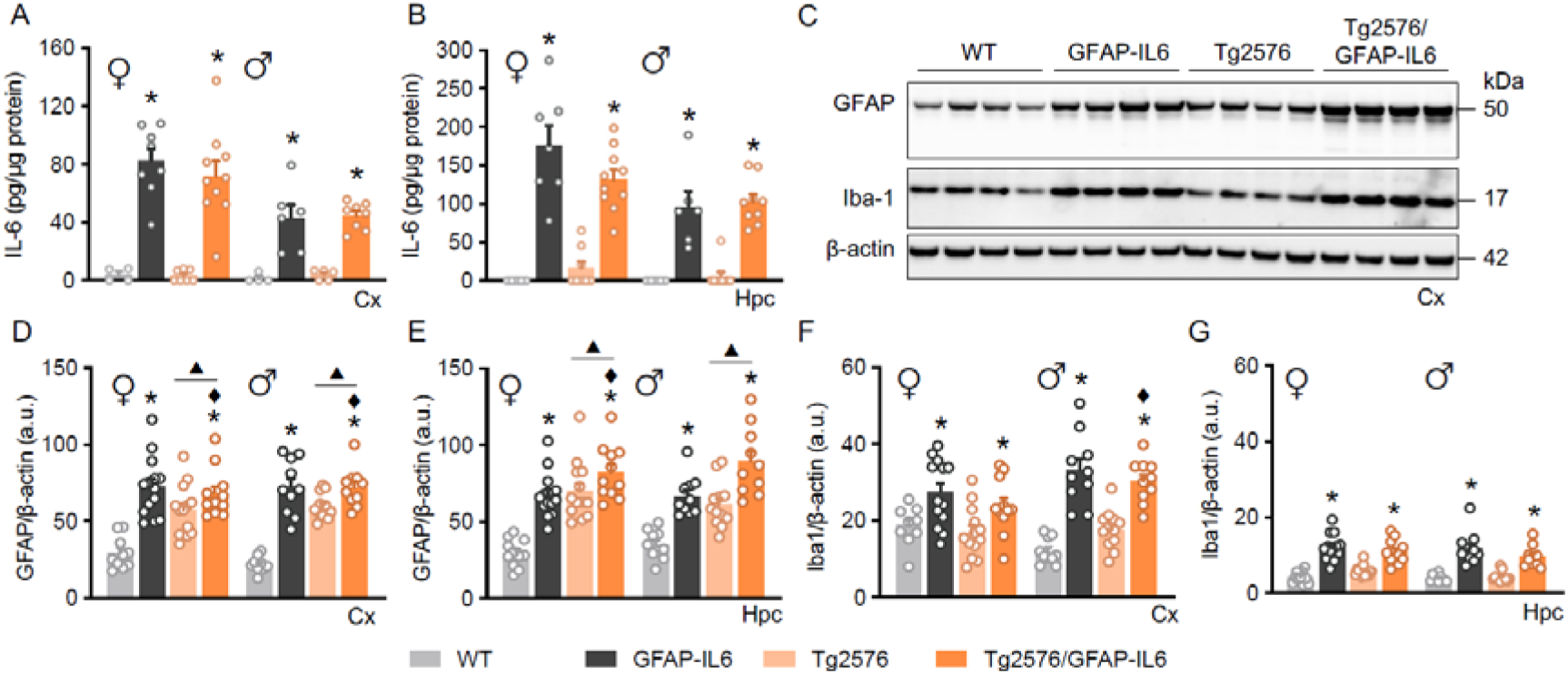
Analysis of cortical and hippocampal gliosis markers. **A,B** Cortical and hippocampal IL-6 levels. **C** Representative cortical GFAP and Iba1 immunoblots. **D - G** Quantification of GFAP (D, E) and Iba1 (F, G) levels relative to β-actin in the cortex and hippocampus. Data are mean ± SEM; analyzed by GzLM statistical test; ▴ p ≤ .05 Tg2576 effect, **\*** - p ≤ .05 interaction between both factors. Cx, cortex; Hpc, hippocampus.

Whereas the astrocytic marker GFAP was elevated in cortical and hippocampal homogenates of Tg2576 and Tg2576/GFAP-IL6 mice, the microglial marker Iba1 was not affected in these mice. Importantly, increased levels of both markers were observed in the cortex and hippocampus in response to central-targeted IL-6 overexpression in both GFAP-IL6 and Tg2576/GFAP-IL6 mice (Fig. 5C–G, Supplementary Fig. 2).

Similar results were observed by immunostaining the cortex and hippocampus of these mice (Fig. 6A, B, D, E) which uncover the main effect of central-targeted IL-6 in astrogliosis and microgliosis not only in basal but also in pathological conditions.

**Figure 6.**
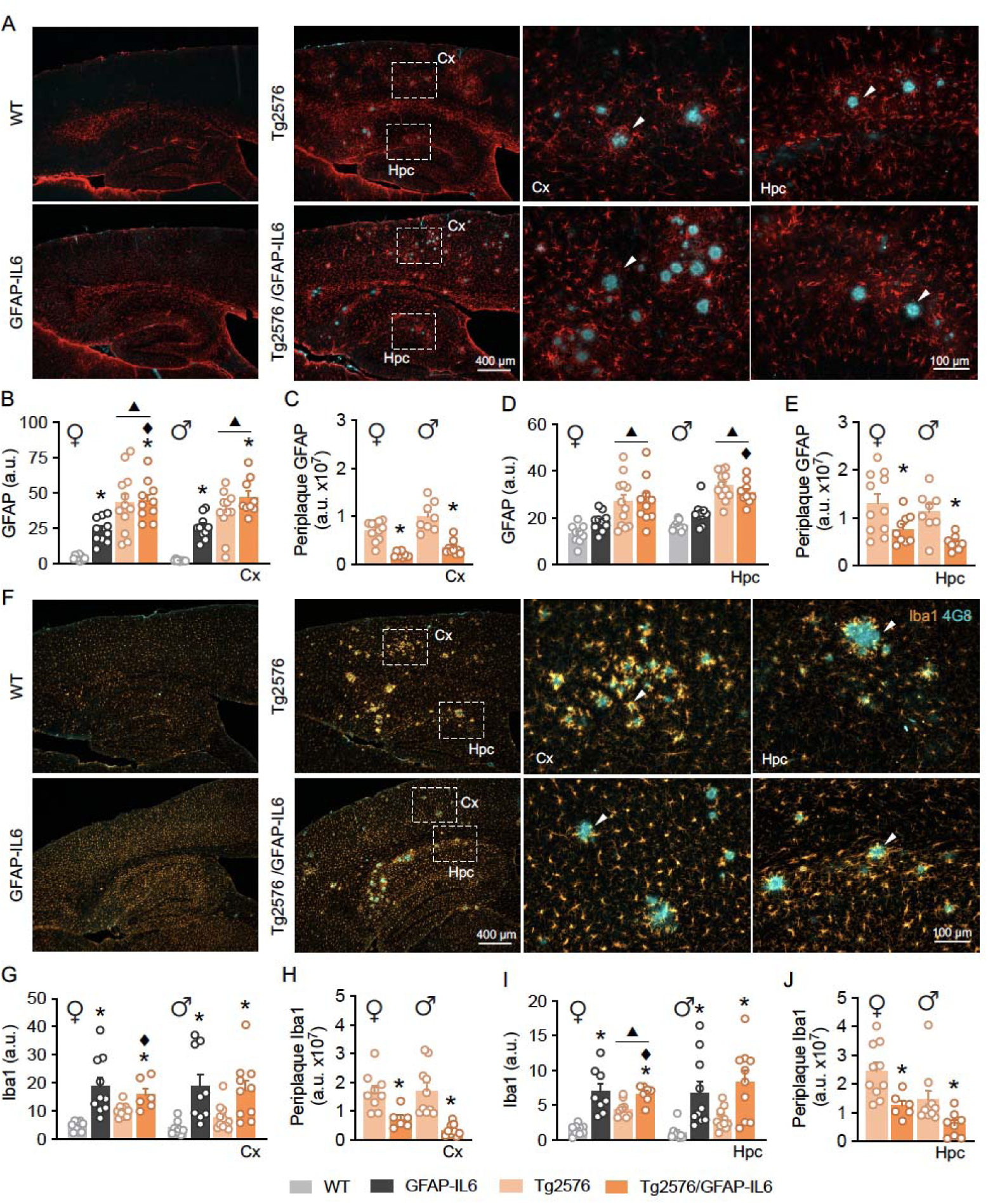
Distribution of glial markers surrounding cortical and hippocampal amyloid plaques in Tg2576 mice. **A** Representative GFAP and 4G8 immunostaining, higher cortical *(middle)* and hippocampal *(right)* magnifications. White arrows indicate glial cells surrounding amyloid plaques. **B, D** Quantification of GFAP levels in the cortex and hippocampus. **C, E** Percentage of GFAP surrounding cortical and hippocampal amyloid plaques. **F** Representative Iba1 and 4G8 immunostaining, higher cortical *(middle)* and hippocampal *(right)* magnifications. White arrows indicate glial cells surrounding amyloid plaques. **G, I** Quantification of Iba1 levels in the cortex and hippocampus. **H, J** Percentage of Iba1 surrounding cortical and hippocampal amyloid plaques. Data are mean ± SEM; analyzed by GzLM statistical test. ▴ p ≤ .05 Tg2576 effect, **\*** - p ≤ .05 interaction between both factors. Cx, cortex; Hpc, hippocampus.

Finally, we analyzed the glial reactivity specifically in the vicinity of amyloid plaques, which highlighted distinct patterns in Tg2576 and Tg2576/GFAP-IL6 mice. Astrocytosis and microgliosis were predominantly localized around amyloid plaques in Tg2576 mice, while this periplaque reactivity was markedly reduced in Tg2576/GFAP-IL6 animals (Fig. 6C, E, H, J), which may underpin a differential distribution of astrocytes and microglia both in cortex and hippocampus in pathological conditions in response to central-targeted IL-6 overexpression.

### 2.5. Chronic central-targeted IL-6 overexpression upregulates cortical immune activation, phagocytic signaling, and gliosis in aged Tg2576 mice

In humans, elevated IL-6 levels are associated with reduced volume in both subcortical regions, such as the hippocampus, and cortical areas, suggesting that IL-6 may play a critical role in regulating these areas [40]. Therefore, we conducted an unbiased transcriptomic analysis of aged male and female WT, GFAP-IL6, Tg2576, and Tg2576/GFAP-IL6 mice to uncover the cortical signature associated with AD-like disease and sustained by central-targeted IL-6 overexpression (Supplementary Fig. 3A–C). Transcriptomic profiling revealed extensive gene expression changes in Tg2576 mice compared to WT mice, with sex-specific patterns (Fig. 7A-C). A total of 241 differential expressed genes (DEGs) were upregulated in Tg2576 females and 181 in males, with 10 genes commonly upregulated in both sexes (Fig. 7A). Heatmap and volcano plot analyses highlighted distinct transcriptional signatures, showing robust upregulation of glial reactivity and neuroinflammatory genes, including astrocytic *Gfap* and microglial *Tyrobp*, *Lyz2*, and *Mrc1*. Male-specific microglial-associated genes included *Fcgr3* and *Mpeg1*, while *Clec7a* was elevated in females. Females also displayed upregulation of genes linked to amyloid interaction (*Zic1*, *Col25a1*), synaptic integrity (*Ptpro*, *Cpeb2*, *Amigo2*), oxidative stress response (*Ngb*), and cytokine signaling (*Dlx1*), whereas neuronal maintenance transcripts (*Pla2g4e*, *Kif5a*) and APP-processing genes (*Adamtsl4*, *Scg5*) were downregulated (Fig 7B, C).

**Figure 7.**
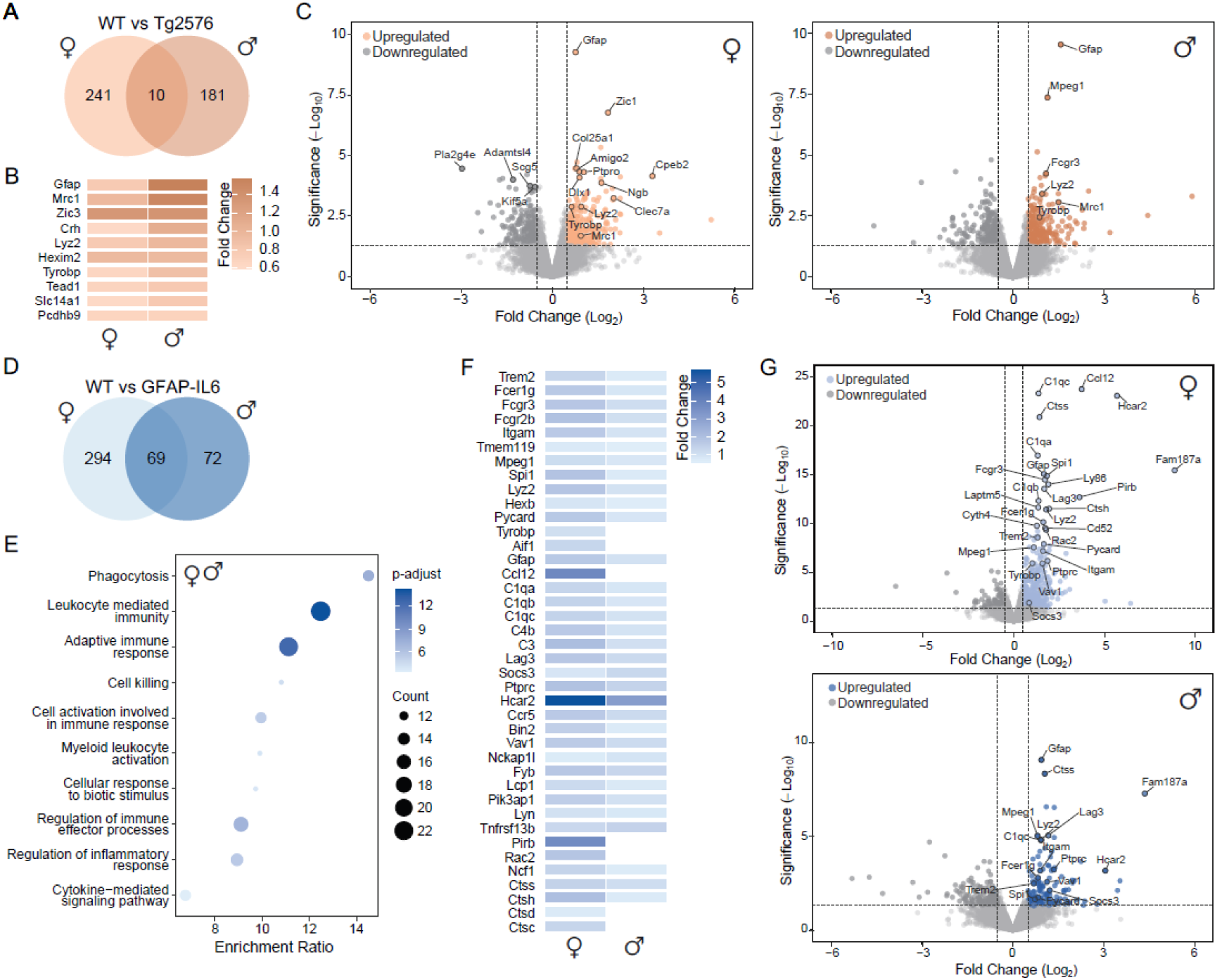
Transcriptomic comparisons and functional enrichment in Tg2576 and GFAP-IL6 mice compared to WT mice. **A-C** WT vs Tg2576 analysis. Venn diagram illustrates the overlap of upregulated genes in female and male mice (A). Heatmap of relative fold changes for shared genes upregulated in both sexes (B). Volcano plots of significantly upregulated DEGs in female and male mice (C). **D - G** WT vs GFAP-IL6 analysis. Venn diagram comparing upregulated genes in female and male mice (D). Dot plot visualizes the top enriched GO Biological Process terms for the shared upregulated genes (E). Heatmap of genes overrepresented in top GO terms in both sexes (F). Volcano plots of upregulated genes in females and males (G). All data were derived from bulk cortical transcriptomes. DEGs were defined by adjusted p-value < 0.05 and corresponding log2 fold change thresholds. Heatmaps display log2 fold change; volcano plots ranked genes by statistical significance (-log10 adjusted p-value) and log2 fold change; dot plots scale by enrichment ratio (log2 fold change), with dot size indicating gene count and color representing enrichment significance. DEG, Differentially Expressed Gene; GO, Gene Ontology; FC, Fold Change.

When GFAP-IL6 mice was compared to WT mice, a robust upregulation of immune and inflammatory pathways was observed in both sexes (Fig. 7D-G). A total of 294 and 72 DEGs were upregulated in GFAP-IL6 females and males, respectively, with an additional 69 genes commonly upregulated in both sexes (Fig. 7D). Gene Ontology (GO) analysis revealed that these shared DEGs are enriched for biological processes related to phagocytosis, leukocyte-mediated immunity, adaptive immune response, and cell killing, among others (Fig. 7E), with shared genes within the overrepresented GO terms (Fig. 7F). Key overlapping genes included microglial regulators (*Trem2*, *Fcer1g*, *Fcgr3*, *Fcgr2b*, *Itgam*, *Tmem119*, *Mpeg1*, *Spi1*, *Lyz2*, *Hexb*, and *Pycard*) and astrocytic *Gfap*. Additional immune genes such as *Fam187a*, along with complement components (*C1qa*, *C1qb*, *C1qc*, *C4b*, *C3*), immune checkpoints/regulatory factors (*Lag3*, *Socs3*, *Ptprc*), and pro-inflammatory and phagocytosis-associated pathways (*Hcar2*, *Ccr5*, *Bin2*, *Vav1*, *Nckap1l*, *Fyb*, *Lcp1*, *Pik3ap1*, *Lyn*, *Tnfrsf13b*) were strongly upregulated. Female-specific transcriptional upregulation was noted for *Tyrobp*, *Aif1*, *Ccl12*, *Pirb*, and *Rac2*, accompanied by enhanced expression of lysosomal proteases (*Ncf1*, *Ctss*, *Ctsh*, *Ctsd*, *Ctsc*) (Fig. 7G).

Comparison of Tg2576/GFAP-IL6 double transgenic mice with Tg2576 mice revealed an upregulation of genes associated with immune activation, phagocytic signaling, and gliosis (Fig. 8A-G). A total of 227 genes were upregulated in Tg2576/GFAP-IL6 females and 285 in males, with 94 shared DEGs primarily enriched for GO categories including antigen processing and presentation, leukocyte mediated immunity, cell killing, phagocytosis, adaptive immune response, and regulation of immune effector processes, among others. Commonly upregulated genes include regulators of microglial activation and phagocytosis (*Trem2*, *Spi1*, *Fcgr2b*, *Fcgr3*, *Fcer1g*, *Itgam*, *Aif1*, *Lag3*, *Irf8*, *Ly86*, *Ptprc*, *Nckap1l*, *Rac2*, *Vav1*, *Lcp1*, *Arpc1b*, *Bin2*, *Pik3ap1*, *Cd300c2*), complement cascade components (*C1qa*, *C1qb*, *C1qc*, *C3*, *C4b*), and lysosomal or proteolytic enzymes (*Ctss*, *Ctsh*, *Ctsc*, *Lyz2*, *Pycard*). Glial reactivity markers (*Gfap*, *Tspo*, *Il4ra*, *Socs3*, *Csf2rb*), integrins and extracellular matrix–related genes (*Itgb2*, *Lrg1*, *Cfh*), and cytokine or chemokine-related transcripts (*Ccl12*, *Tnfrsf13b*) were notably upregulated (Fig. 8A-C, F, G). Sex-specific upregulated DEGs in both females and males mapped to similar immune-related GO categories, including cytokine production and broader immune responses (Fig. 8D, E).

**Figure 8.**
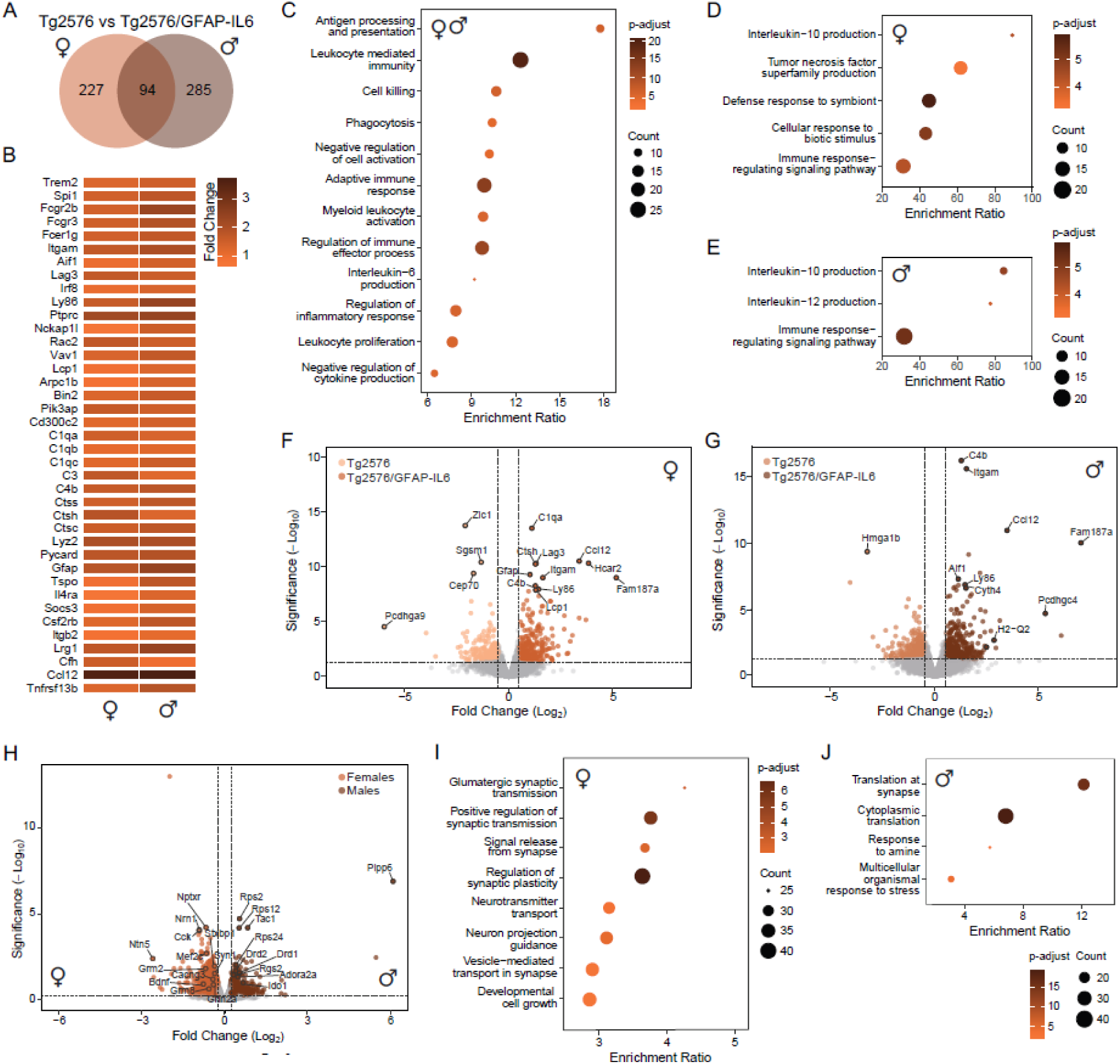
Transcriptomic comparisons and functional enrichment in Tg2576/GFAP-IL6 mice compared to Tg2576 mice. **A–G** Tg2675 vs Tg2576/GFAP-IL6 analysis. Venn diagram for upregulated gene overlap in female and male mice (A). Heatmap of selected shared DEGs in both female and male mice (B). Dot plot of top GO Biological Process categories among shared upregulated genes (C). Dot plots of top GO Biological Process categories linked with overexpressed genes in females (D) and males (E). Volcano plots of significantly upregulated DEGs between Tg2576 and Tg2576/GFAP-IL6 females (F) and males (G), with relevant genes annotated. **H** Volcano plot of significantly upregulated DEGs between Tg2576/GFAP-IL6 females and males, with relevant genes annotated. **I - J** Dot plot of enriched GO Biological Process categories in DEGs in females (I) and males (J). Data represent transcriptomes derived from bulk Cx tissue, DEGs are defined by adjusted p-value < 0.05 and log2 fold change thresholds. Heatmaps are represented by log2FC, volcano plots are ranked by statistical significance (-log10padj) and log2FC, dot plots are scaled by log2FC (enrichment ratio), with dot size indicating gene count and color representing enrichment significance. DEG, Differentially Expressed Gene; GO, Gene Ontology; FC, Fold Change.

Analyses of sex differences focusing on upregulated genes in Tg2576/GFAP-IL6 females versus males revealed distinct functional signatures, with females exhibiting enrichment primarily in synaptic and neurotransmission-associated pathways, whereas males showed strong activation of translational and cellular stress responses. In females, relevant genes among those top GO categories included multiple glutamate receptor subunits (*Grin1*, *Grin2a*, *Grin2b*, *Grm2*, *Grm8*), calcium and synaptic vesicle regulators (*Cacng2*, *Cacng3*, *Stxbp1*, *Syt1*, *Syn1*, *Syn2*, *Snap47*, *Unc13a*, *Unc13b*), neural development and synaptic plasticity genes (*Nrn1*, *Ntn5*), and modulators of neuronal growth and guidance (*Bdnf*, *Epha4*, *Sema3c*, *Sema7a*, *Plxna1*, *Hdac6*, *Mef2c*). In males, upregulated genes comprised several ribosomal proteins (*Rpl12*, *Rpl13a*, *Rpl35*, *Rpl38*, *Rps2*, *Rps12*, *Rps11*, *Rps24*, *Rps27a*, *Rps28*), dopamine and adenosine receptors (*Drd1*, *Drd2*, *Adora2a*), and signaling modulators (*Rgs2*, *Rgs9*, *Tac1*, *Ntrk1*, *Penk*, *Ido1*) (Fig. 8H-J). A t-SNE plot was used to visualize the dispersion of GO biological processes across the experimental comparisons (Tg2575 vs Tg2576/GFAP-IL6, WT vs Tg2576, and WT vs Tg2576/GFAP-IL6) in each sex and revealed that principal categories consistently clustered in all groups were related to RNA processing and autophagy (Supplementary Fig. 3D, E).

## 3. Discussion

Here we reveal that chronic central-targeted IL-6 overexpression significantly contributed to the molecular landscape underlying amyloidosis, increasing cortical and hippocampal Aβ_42_/Aβ_40_ ratios and gliosis outside of plaques in aged Tg2576 males and females.

Neuroinflammation is a central mechanism in Alzheimer’s disease, with increasing evidence positioning IL-6 as a key mediator of disease initiation and progression [8–10,41]. A recent study has highlighted that IL-6 deficiency improves the cognition, reduces Aβ plaque accumulation and mitigates AD-like associated hippocampal neuroinflammation by inhibiting STAT3-cGAS-STING pathway [42]. Based on our results showing that central-targeted IL-6 chronic overexpression significantly influences physiological, behavioral, and hippocampal and cortical molecular features in the Tg2576 AD mouse model, this reveals the importance of a central source of IL-6 on this disease.

Premature mortality was evident in both male and female Tg2576 mice. This phenomenon may be partly attributable to Aβ toxicity, consistent with previous reports showing that reducing Aβ production or enhancing its clearance reduces mortality in mice [43]. However, the mortality associated with APP overexpression may also stem from Aβ-independent factors, including seizures, cerebrovascular dysfunction, oxidative stress, genetic background, environmental variables, and sex-specific mechanisms [24,44–47]. Although AD-induced mortality was reversed by the inhibition of astrocytic IL-6 trans-signaling [17], central-targeted IL-6 overexpression did not impact on survival. This emphasizes the inhibition of astrocytic IL-6 trans-signaling as a key pathway mediating the neuroinflammatory response to damage, partially rescuing the premature mortality in Tg2576 mice [17]. IL-6’s influence on blood-brain barrier maintenance and neuronal signaling likely contributes to this increased vulnerability [48]. Weight loss, a clinical hallmark of AD observed in both human patients and transgenic mouse models [17,49,50], was evident in Tg2576 mice and further exacerbated by central-targeted IL-6 overexpression in females. Notably, central-targeted IL-6 overexpressing mice, both wild-type and Tg2576, exhibited reduced body weight in females, as well as reduced inguinal white fat depots in both males and females, consistent with the cytokine’s established role in the central regulation of body weight, energy homeostasis, and adiposity [51,52]. These findings suggest that central-targeted IL-6 contributes to both physiological and AD-associated metabolic decline.

Behavioral assessment confirmed the early-onset and progressive behavioral alterations in Tg2576 mice, consistent with previous studies [18,20,21,53–56]. Behavior was modulated by IL-6 in an age-and sex-dependent manner, potentiating hyperactivity and exploration in young Tg2576 mice, while dampening exploration and locomotion in aged animals. These findings align with earlier reports describing hypoactivity and reduced exploration in IL-6-deficient mice [25,57,58], despite some conflicting results [59]. Additionally, an anxiolytic-like role of IL-6 has been previously described [60]. Consistent with this, central-targeted IL-6 overexpression in our study further reduced anxiety-like behaviors in both young and aged Tg2576 mice. Conversely, increased anxiety has been observed in IL-6-deficient models [25,57,61]. Cognitive analyses revealed Tg2576-associated deficits in working and spatial memory, accompanied by sex-dependent differences in performance, potentially reflecting task-specific sensitivity, as previously reported [20]. These cognitive impairments have been linked to deficits in hippocampal long-term potentiation (LTP) [53,62]. IL-6 is known to negatively influence hippocampal-dependent cognition [63,64], whereas inhibition of endogenous IL-6 in healthy brains enhances LTP and improves performance in hippocampal-dependent memory tasks [14,65]. In our study, central-targeted IL-6 overexpression selectively improved working memory while impairing spatial reference memory. Similar age-dependent effects of IL-6 on cognition have been reported in both IL-6-deficient [66,67] and IL-6-overexpressing mice [68], potentially driven by age-related neuropathology, reduced LTP, and cerebellar atrophy [27,69]. Collectively, these findings suggest that IL-6 exerts a context-dependent role in cognitive function, modulated by age, sex, brain region, and cognitive domain. Disruptions in IL-6 signaling may therefore alter brain development and, consequently, brain functions [39,65,66]. Motor coordination deficits, a recognized late-stage feature of AD [70], were evident in Tg2576 mice, consistent with cortical dysfunction and Aβ-induced impairments [20,54,71,72]. GFAP-IL6 mice also exhibited progressive neurological dysfunction, including ataxia, motor impairment, seizures, and premature mortality [22,27,73], and while the effect of IL-6 on motor performance in Tg2576 mice was modest, it likely contributes to overall functional decline.

At the molecular level, mutant APP overexpression in Tg2576 mice results in increased secretion of amyloid beta peptides, specifically Aβ_42_ and Aβ_40_, leading to the formation of senile plaques with dense amyloid cores in the cortex and hippocampus [18], which was confirmed in our study. Inflammatory mediators as IL-6 have been reported to enhance APP expression and processing and influence Aβ deposition, potentially establishing a positive feedback loop that contributes to neurodegeneration [74,75]. Whereas central-targeted IL-6 overexpression did not impact either hippocampal or cortical APP, it decreased total soluble Aβ peptide levels in both brain areas. Notably, central-targeted IL-6 overexpression increased the cortical and hippocampal Aβ_42_/Aβ_40_ ratio, suggesting selective upregulation of the more pathogenic Aβ_42_ isoform and promoting plaque formation. This contrasted with reports from other AD models [12,42], pointing to model- and context-specific effects.

Glial reactivity, typified by astrogliosis and microgliosis, was robust in Tg2576 mice, as previously reported [3,4]. Glial reactivity is also a common hallmark of GFAP-IL6 mice, which develop a progressive neuroinflammatory state in the brain parenchyma due to the chronic production of IL-6 in the brain [64]. While gliosis was predominantly associated with amyloid plaques in Tg2576 mice, central-targeted IL-6 overexpression dramatically decreased the glial reactivity responding to amyloid plaques. This strongly suggests an alteration in the physiological responses of glial cells to amyloid plaques in GFAP-IL6 mice. Thus, the moderate increase in amyloid plaque load observed in IL-6-overexpressing mice could be attributed to a potential overload of glial cell phagocytic capacity, resulting in ineffective clearance. The microenvironment surrounding amyloid plaques appears to be crucial in modulating glial cell phagocytic function and influencing Aβ deposit clearance. Indeed, the heterogeneous nature of microglial responses in the context of plaque formation has already been reported by other studies [4,76]. Finally, it is noteworthy that GFAP-IL6 mice display glial cell reactivity at early stages in mice [64], preceding plaque formation, which could potentially affect the pattern observed in our mice.

The cortical transcriptomic profiling revealed extensive gene expression alterations across Tg2576, GFAP-IL6, and Tg2576/GFAP-IL6 mice, highlighting distinct yet convergent neuroinflammatory and synaptic signatures associated with Alzheimer’s disease-like pathology [77]. In Tg2576 mice, upregulation of *Gfap* confirmed robust astrocyte reactivity, consistent with the reactive gliosis characteristic of this model [3]. Enhanced expression of *Mpeg1* and *Fcgr3*, associated with activated and plaque-encapsulating microglia in AD [78–80], reflected elevated immune responses to amyloid pathology. Increased transcription factors *Zic1*, which can transactivate *Apoe* [81], and *Dlx1*, a regulator of neuronal differentiation [82], suggest disrupted neuronal development and amyloid metabolism. Elevated *Col25a1*, known to bind amyloid beta peptides, further supports the promotion of plaque formation observed in AD models [83] and human tissue [84]. Upregulation of *Amigo2*, *Ptpro*, and *Cpeb2* suggests modulation of synaptic maintenance and signal transduction, with *Amigo2* potentially linked to reactive astrocyte responses [85], while elevated *Ngb* hints activation of antioxidative mechanisms in response to Aβ-induced mitochondrial stress [86]. Conversely, downregulation of *Pla2g4e* and *Kif5a*, genes related to membrane trafficking and axonal transport, is consistent with synaptic dysfunction induced by amyloid accumulation, in agreement with previous findings linking their loss to synaptic and mitochondrial vulnerability [87,88]. Collectively, Tg2576 mice exhibit transcriptional hallmarks of neuroinflammation, synaptic dysregulation, and neuronal stress reflecting early AD pathology.

GFAP-IL6 mice, in contrast, displayed a transcriptional mapping dominated by immune activation and glial signaling, driven by chronic central-targeted IL-6 overexpression. Upregulated microglial receptors, microglial markers and immune regulators (*Trem2*, *Tyrobp*, *Fcgr2b*, *Fcgr3*, *Fcgr4*, *Ccr5*, *Cx3cr1, Lag3, Pirb, Spi1, Aif1, Tmem119, Itgam*) underscore enhanced microglial activation and leukocyte-like immune surveillance already described in this mouse model [22,27,68,89]. Upregulation of lysosomal and proteolytic genes (*Ctss*, *Ctsh*, *Ctsd*, *Hexb*) suggests extensive microglial reactivity and phagolysosomal remodeling [90,91], while increased cytoskeletal regulators (*Rac2*, *Vav1*, *Bin2*) indicate augmented glial motility and morphological transformation. These signatures, alongside complement activation (*C1qa-c*, *C3*, *C4b*), suggest an environment primed for phagocytosis, antigen presentation, and adaptive immune engagement already described in this model of neuroinflammation [22,89,92–95]. Interestingly, upregulation of *Hcar2* (GPR109A), known for mediating anti-inflammatory responses and metabolic regulation by suppressing fat breakdown [96,97], links the neuroinflammatory activation with the altered metabolic states observed in GFAP-IL6 mice. The concurrent upregulation of fatty acid synthesis regulators (*Cebpa*, *Cebpb*), which promote fat cell formation and storage [98], and of metabolic genes (*Ucp2*, *Ltc4s*, *Hpgds*), together with reduced *Hmgs2* expression, involved in fat utilization during fasting [99], suggests a metabolic shift toward decreased lipid metabolism and increased fat accumulation.

The Tg2576/GFAP-IL6 double transgenic mice exhibited a pronounced cortical activation of the transcriptional signature. When compared to Tg2576 mice, enriched pathways indicated a shift toward increased immune cell recruitment, maturation, and a potential adaptive immune landscape, and showed a synergistic upregulation of genes involved in phagocytosis, cell killing, and cytotoxicity. Most shared DEGs are common with those upregulated in GFAP-IL6 mice, including major microglial regulators, astrocytic markers, complement cascade components, and lysosomal enzymes, indicating a potent inflammatory milieu combining amyloid-dependent and cytokine-driven immune activation. This pattern aligns with current views that inflammation in the AD brain critically shapes the disease trajectory, rather than acting as an isolated trigger. Consistent with this notion, IL-6-driven immune activation could have a dual role, being initially protective by enhancing clearance but becoming detrimental when chronically sustained [100–103]. Interestingly, different transcriptional profiles are revealed in males and females. Females displayed enrichment of pathways related to synaptic signaling, neurotransmitter transport, vesicle cycling, and neuronal projection guidance. These included glutamatergic transmission and plasticity networks (*Grin1*, *Grin2a*, *Grin2b*, *Grm2*, *Adcyap1*, *Mef2c*, *Bdnf*, *Stxbp1*, *Syn1*, *Stx1a*), which could be related to attempts to preserve synaptic integrity under chronic inflammatory stress [104–107]. Conversely, male mice exhibited gene enrichment centered on local translation and cellular stress adaptation, including ribosomal gene clusters (*Rpl12*, *Rpl13a*, *Rps5*, *Rps27a*) and transcripts implicated in amine and dopaminergic signaling (*Drd1*, *Drd2*, *Adora2a*, *Pde1b*, *Rgs9*, *Th*) [108,109]. These data suggest that while females engage synaptic remodeling programs to counteract inflammation-induced synaptic vulnerability, males prioritize translational and metabolic adaptations suited for stress management and protein turnover.

From a translational perspective, our findings suggest that central-targeted IL-6 is an important source to trigger AD pathophysiology. However, the translatability of these preclinical findings is limited, since the present study was not conducted at thermoneutrality and use genetic mice. Moreover, Tg2576 mice are characterized by Aβ deposition, but in most AD patients, tau pathology is also implicated [110,111]. It is important to note that although GFAP is expressed mainly in astrocytes, some hippocampal neural progenitors in the mouse adult also display *Gfap* expression, which may contribute the phenotype in GFAP-IL6 and Tg2576/GFAP-IL6 [112].

In conclusion, our findings emphasize the multifaceted nature of central-targeted IL-6 in mediating AD physiopathology in Tg2576 males and females. Given its implication from early stages and its pleiotropic effects, targeting this cytokine may represent a promising strategy to manage symptoms in the initial phases of the disease.

## Lead Contact

Further information and requests for resources and reagents should be directed to Paula Sanchis and Juan Hidalgo.

## Materials Availability

This study did not generate new unique materials.

## Supplementary data

Supplementary Figures 1-3

## Data and Code Availability

The transcriptomic raw RNA-seq data have been deposited at the Sequence Read Archive Database (https://www.ncbi.nlm.nih.gov/sra/) under BioProject accession PRJNA1369206.

## Consent for publication

Not applicable.

## Authors’ contributions

C.C. and J.H. conceived and designed the study. C.C., P.S, K.A., G.C., M.G., and E.S. performed research. C.C. performed the formal analysis of the data. C.C. P.S. and J.H. wrote the initial draft of the manuscript. All authors edited and reviewed the manuscript. The authors claim no competing interests.

## Acknowledgements

The GFAP-IL6 mice were obtained from The Scripps Research Institute (La Jolla, USA) where they were developed by I. L. Campbell (28). C.C. and K.A. acknowledge the support of PIF UAB and FPU17/02065 fellowships, respectively. P.S. received the Lundbeck R380-2021-1300 fellowship.

