## Supplementary figures and images for "Chronic central-targeted Interleukin-6 overexpression promotes hippocampal and cortical neuropathology in the Tg2576 mouse model of Alzheimer’s disease"

### Supplemental Figure 1

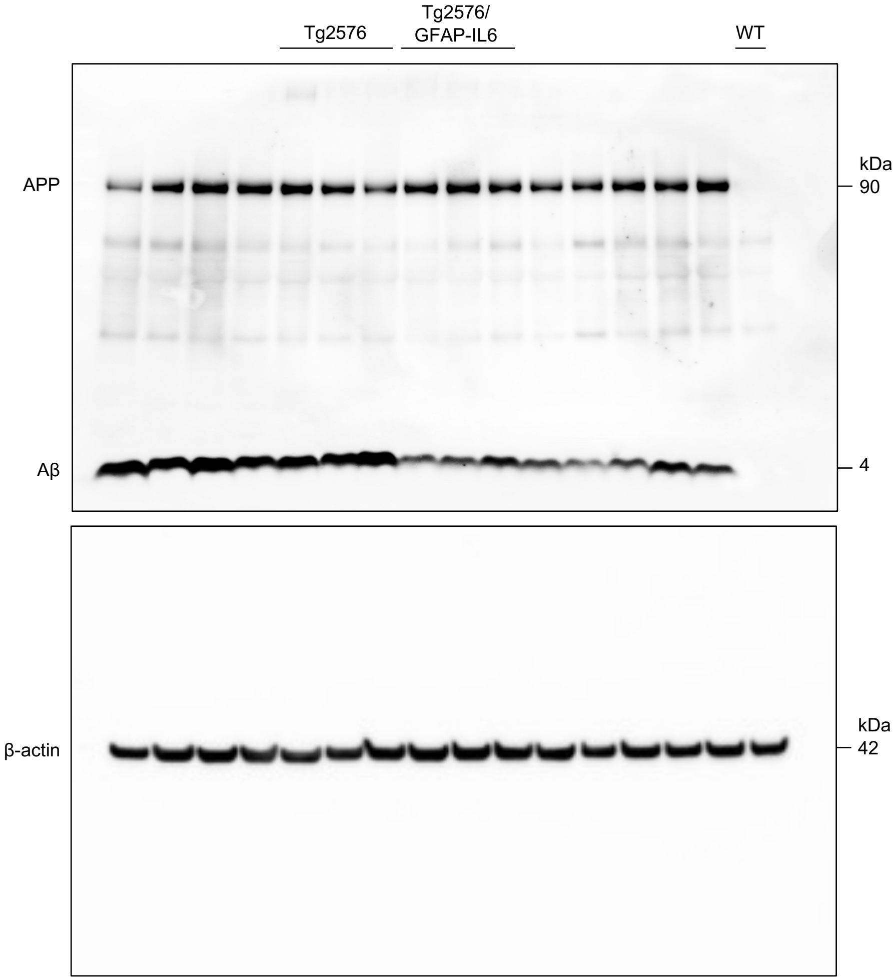

### Supplemental Figure 2

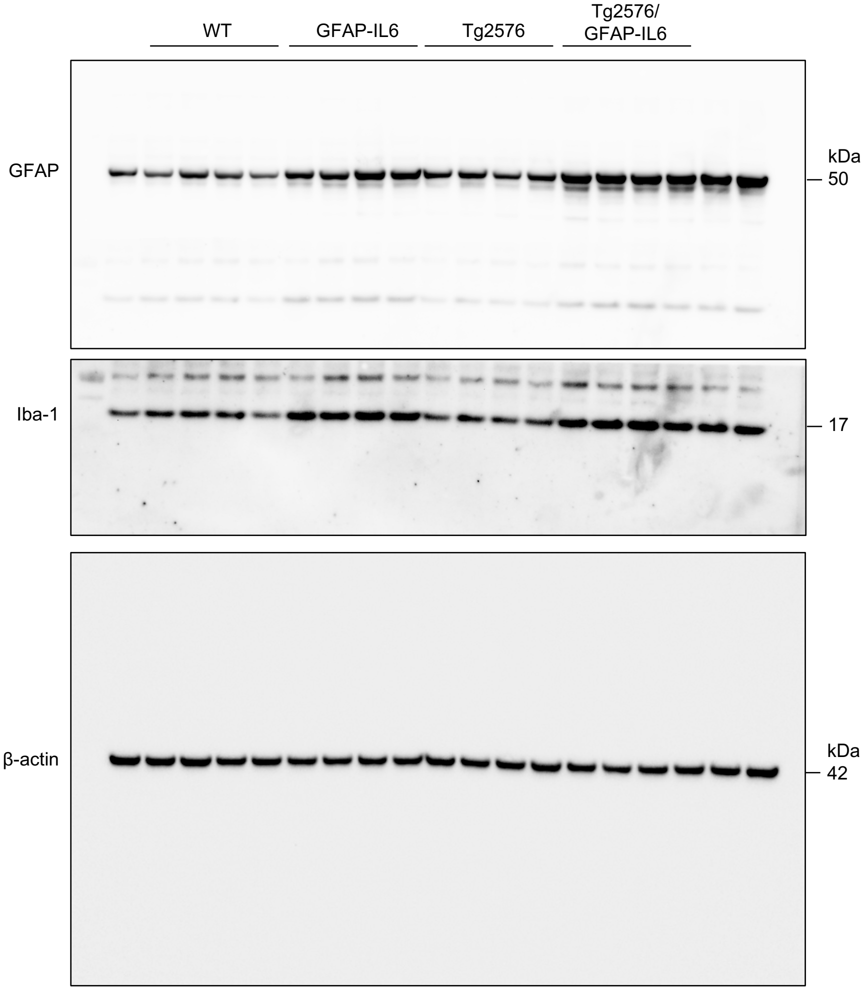
